# Targeting staphylococcal cell-wall biosynthesis protein FemX through steered molecular dynamics and drug-repurposing approach

**DOI:** 10.1101/2023.06.13.544722

**Authors:** Shakilur Rahman, Subham Nath, Utpal Mohan, Amit Kumar Das

## Abstract

*Staphylococcus aureus*-mediated infection is a serious threat in this antimicrobial-resistant world. *S. aureus* has become a ‘superbug’ by challenging conventional as well as modern treatment strategies. Nowadays, drug repurposing has become a new trend for the discovery of new drug molecules. This study focuses on evaluating FDA-approved drugs that can be repurposed against *S. aureus* infection. Steered molecular dynamics (SMD) has been performed for Lumacaftor and Olaparib against staphylococcal FemX to understand their binding to the active site. A time-dependent external force or rupture has been applied to the ligands to calculate the force required to dislocate the ligand from the binding pocket. SMD analysis indicates that Lumacaftor has a high affinity for the substrate binding pocket in comparison to Olaparib. Umbrella sampling exhibits that Lumacaftor possesses a higher free energy barrier to displace it from the ligand-binding site. The bactericidal activity of Lumacaftor and Olaparib has been tested, and it shows that Lumacaftor has shown moderate activity along with biofilm inhibition potential (MIC value with conc. 128 μg/mL). Pharmacokinetic and toxicology evaluations indicate that Lumacaftor has higher pharmacokinetic potential with lower toxicity. This is the first experimental report where staphylococcal FemX has been targeted for the discovery of new drugs. It is suggested that Lumacaftor may be a potential lead molecule against *S. aureus*.

## 1. Introduction

The increasing prevalence of methicillin-resistance *Staphylococcus aureus* (MRSA) infections has gained attention to design and develop novel antibacterial agents^1^. *Staphylococcus aureus* is one of the deadly microorganisms from the ESKAPE family of pathogens^2^. The resistance strategy of *S. aureus* to methicillin and other β-lactam antibiotics is generated by expressing a foreign penicillin-binding protein (PBP) known as PBP2a^3^. PBP2a can perform the functions of host PBPs and bypass the action of β-lactam antibiotics. Additionally, several other auxillary (aux) genes have also been identified that play a crucial role in methicillin resistance, namely, **f**actors **<u>e</u>**ssential for **<u>m</u>**ethicillin-resistance (fem) family genes^4^. Three of these genes, *femX, femA* and *femB*, play a vital role in the latter stage of the peptidoglycan biosynthesis process of *Staphylococci* and can be served as potential targets for developing new antimicrobial therapeutics^5,6^. Staphylococcal peptidoglycan (PG)-repeat unit consists of a disaccharide, a pentapeptide stem and a penta-glycine bridge structure. The peptidoglycan disaccharide unit (DU) is composed of *N*-acetyl-glucosamine (NAG) and *N*-acetyl-muramic acid (NAM) and is conserved among all eubacteria whereas the pentapeptide stem and bridge structure vary between species^7^. FemX, FemA and FemB add five glycine residues to the lysine of the A-E-K-A-A pentapeptide stem, thereby developing the flanking bridge structure. FemX adds the first glycine, FemA adds the second and third glycine, and FemB adds the fourth and fifth glycine to the lysine of the pentapeptide stem^8^.

This pentaglycine (Gly_5_) bridge is essential for crosslinking several PG chains to develop a 20-40 nm thick cell wall ^9,10^. The C-terminal of the Gly_5_ bridge forms an amide bond to the side chain nitrogen of the L-lysine of the A-E-K-A-A pentapeptide stem ^11^. During the final stage of PG synthesis, a mature cell wall was developed by crosslinking the N-terminal of a pentaglycine bridge structure to the D-Ala (4^th^ amino acid of a pentapeptide stem) of a neighbouring PG chain with a peptide bond (Figure 1) ^12^. This crosslinking provides the Gram-positive characteristic mesh-like PG structure to the cell wall. All these fem family enzymatic reactions are highly substrate-specific, where mutations or deletions of the genes can lead to decreased resistance to β-lactam compounds and peptidoglycan crosslinking ^13–15^. More importantly, the deletion of *femX* in *S. aureus* is lethal ^14,16^, whereas the deletion of *femA* and *femB* results in the synthesis of mono and tri-glycyl segments in bridge structure (Figure 2) ^8,17^.

**Figure 1.**
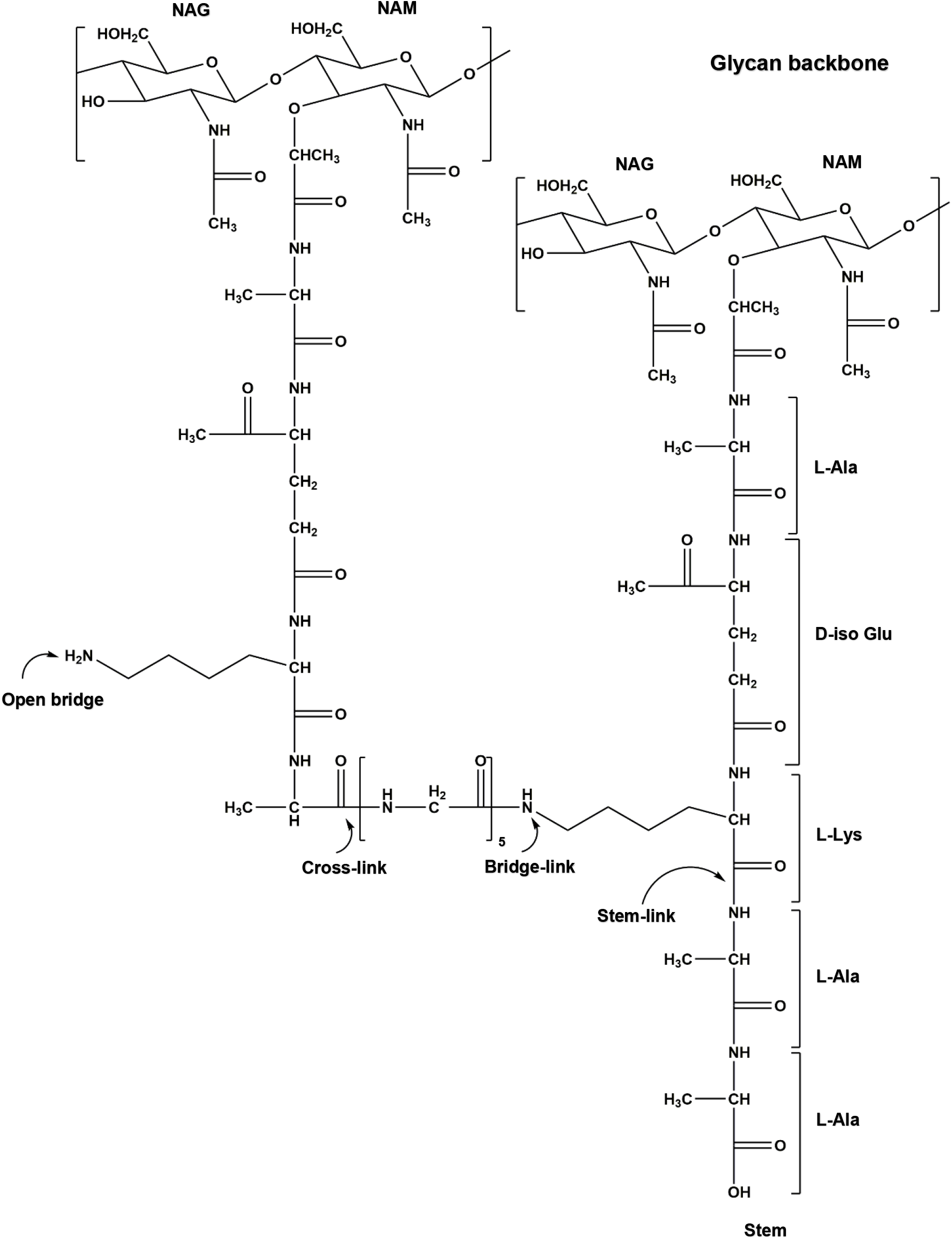
Chemical composition of *S. aureus* peptidoglycan. The disaccharide unit comprises *N*-acetyl-glucosamine (NAG) and *N*-acetyl-muramic acid (NAM). The pentapeptide structure is formed with L-Ala—D-Glu—L-Lys—D-Ala—D-Ala, where the glycine bridge is attached to the ε nitrogen of L-Lys of the third position of the pentapeptide stem. Crosslinking occurs between the N-terminus of the last glycine of the bridge with the D-Ala (fourth position) of the adjacent pentapeptide stem.

**Figure 2.**
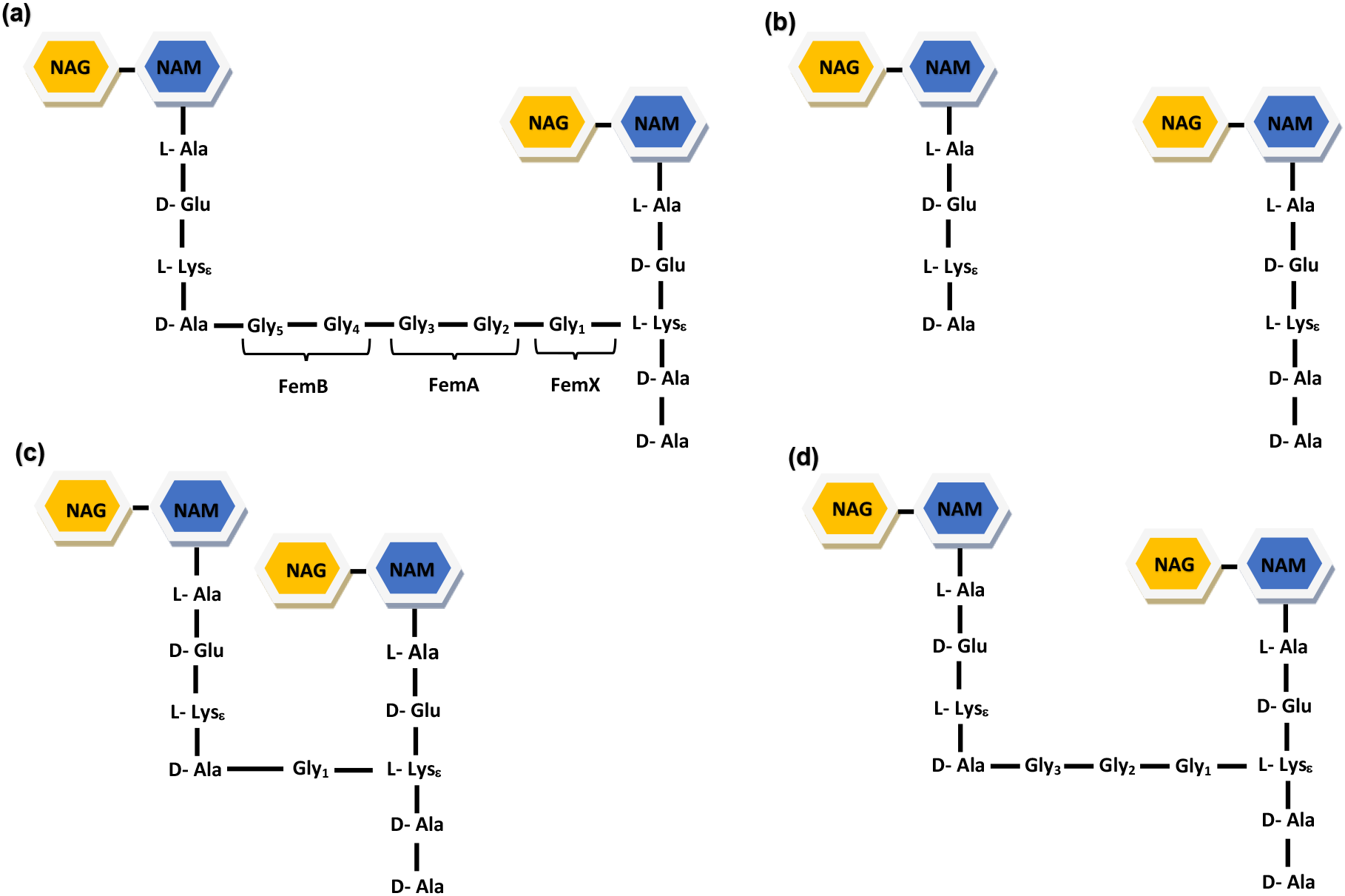
The structure of *S. aureus* peptidoglycan consists of disaccharide, pentapeptide stem (L-Ala—D-Glu—L-Lys—D-Ala—D-Ala) and glycine bridge structure. (a) Proper crosslinking of peptidoglycan chain where mutation/deletion is present either in staphylococcal FemX, FemA or FemB, (b) No crosslinking of peptidoglycan chain due to mutation/deletion in FemX, (c) Very short crosslinking of peptidoglycan chain due to mutation/deletion in FemA, (d) Short crosslinking of peptidoglycan chain due to mutation/deletion in FemB.

In contrast to Gram-negative bacteria, which have an outer membrane, gram-positive bacteria have layers of peptidoglycan that are many times thicker than Gram-negative bacteria. Over the past few decades, the peptidoglycan biosynthesis pathway has been the most attractive target for designing novel antibiotics. PBPs were the primary targets for many years, but due to the accumulation of foreign PBP, treating *S. aureus* has become a severe issue. This has led us to search for new drug molecules against drug-resistant *S. aureus*. The current study presents steered molecular dynamics and umbrella sampling analysis of the FemX-drug complexes, the bactericidal activity of the selected drugs, biofilm inhibition assay and ADMET evaluation of the selected drugs.

## 2. Materials and Methods

### 2.1 Steered dynamics and umbrella sampling

Pulling simulations are essential to estimate the binding energy between Lumacaftor and staphylococcal FemX. The Lumacaftor-FemX and Olaparib-FemX complex structures were obtained from our previous Study ^18^. Protein topology was generated with AMBER99SB forcefield, and ligand topology was generated using the General Amber Force Field (GAFF) ^19^ from the ACPYPE server ^20^. The pulling direction was determined based on the drug-binding pocket of FemX, where the drug-binding pocket was made parallel to the Z-axis. FemX-drug complexes were positioned in a rectangular box with dimensions sufficient to provide a place for pulling simulations to be performed along the Z-axis. The simulation system was solvated using TIP3P water, and counter ions were added to neutralize the net charge of the system. Long-range electrostatic interactions were treated using the particle mesh Ewald (PME) method, and a steepest-descent minimization of 50000 steps was used the remove the bad contacts from the system. The system was equilibrated in an NVT ensemble at 1 atm pressure and 310 K temperature with the Berendsen thermostat. For each system, the FemX backbone was kept constrained, whereas drugs were pulled from the protein active site towards the solvent pull along the Z-axis. Both the ligands were pulled at 0.01 nm/ps by using a spring constant of 1000 kJ mol^-1^ nm^-2^. The final steered molecular dynamics (SMD) simulation of 1000 ps was performed with Berendsen temperature coupling and Parrinello-Rahman pressure coupling using the GROMACS 2020.0 package ^21^. Single pulling vector and exit trajectory were explored during SMD. The snapshots so obtained from SMD trajectories were exploited for umbrella sampling. During sampling, window spacing was kept at 0.1 nm for the centre of mass (COM) separation. The system was further subjected to NPT equilibration for 100 ps. During umbrella sampling (US), a 10 ns simulation was performed for each selected individual configuration. The weighted histogram analysis method (WHAM) module was used to calculate the potential mean force (PMF) from the outcomes of the US simulations.

### 2.2 Bacterial strains, culture conditions and chemical stocks

*Staphylococcus aureus* MTCC 3160 and NCTC 8325 strains (both are methicillin-sensitive; MSSA) were used in this study. A single colony of *S. aureus* was cultured overnight in 5 mL Mueller Hinton broth (MH; HiMedia, India) at 37 °C with 150 rpm agitation. To prepare the stock solution, Lumacaftor and Olaparib (Selleck Chemicals, USA) were dissolved in dimethyl sulfoxide (DMSO; Sigma-Aldrich Inc., USA) to the final concentration of 1mg/mL and stored at -20 °C for further use.

### 2.3 Bactericidal activity assay

Minimum inhibitory concentrations of drugs were calculated based on a previously described broth microdilution method ^22^. 100 μl of Lumacaftor (512 μg/mL) and Olaparib (512 μg/mL) were added individually to the first column of a 96 well-plate (Thermo Fisher Scientific, USA), and two-fold serial dilutions of drugs were added to the other wells. Then, 50 μL of bacterial suspension (10^6^ CFU/mL) was seeded to each well for a final concentration of 10^5^ CFU/mL. Amp-Na (6.25 μg/mL) was used as the positive control. The plate was kept in an incubator at 37 °C for 16 hours. The optical density (OD) of bacterial cultures was measured at 600 nm using a plate reader (BioTek, USA). Each sampling was done in triplicate for quantification. Data are presented as mean standard deviation (SD). Statistical comparisons between groups were performed by Student’s t-test. P< 0.05 was considered to be statistically significant.

### 2.4 Biofilm inhibition assay

A 96-well microtiter plate was prepared according to the method described above. The plate was incubated at 37 °C for 16 hr under static conditions. Trans-chalcone (20 μg/mL) was used as a positive control ^23,24^. After incubation, non-adherent cells were removed, and the wells were washed twice using PBS. The plate was dried for 15 min in laminar airflow, and the biofilm was stained with filtered 0.1% (w/v) crystal violet (CV) for 20 min at room temperature. The excess stain was removed by washing with PBS three times. Then, 95% ethanol was added to the wells and incubated for 10 min. Absorption of the CV stain was recorded at 570 nm using a micro-titre plate reader (BioTek, USA). Each sampling was done in a triplicate manner. Statistical comparisons between groups were performed by Student’s t-test, where p< 0.05 was considered to be statistically significant.

### 2.5 ADME and toxicity evaluation

To evaluate the physicochemical properties and toxicity of the drug, the 2D structure and SMILES of Lumacaftor (PubChem CID: 16678941) and Olaparib (PubChem CID: 23725625) were obtained from the PubChem database ^25^. SwissADME ^26^ was used to calculate the molecular weight, lipophilicity, polarity, insolubility, flexibility, insaturation, GI absorption and blood-brain barrier (BBB) permeation. Further, toxicity properties, such as maximum tolerance dose, cytochrome P450 inhibitory activity, skin sensitization etc., were predicted using the pkCSM tool ^27^.

## 3. Results

### 3.1 Steered dynamics and umbrella sampling

The interaction energy between Lumacaftor and staphylococcal FemX was calculated to understand the strength of the interaction of drug and protein. In steered MD simulations, a time-dependent external force is applied on the ligands to drive its dislocation from the protein, usually which cannot be achieved by standard MD simulation. Notably, the transition between the bound and unbound states has been measured during steered MD. The force was gradually increased concerning time (ps) and distance (nm). Lumacaftor had a steady increase in the applied force until the force reached the maximum value of 3501.17 kJ/mol/nm at around 450ps time, termed rupture force (F_max_). Whereas, Olaparib had a significantly lower F_max_ of 1471.71 kJ/mol/nm at around 200ps time (Figure 3). The force then rapidly decreased and remained constant until the end of the simulation suggesting the disruption in ligand-receptor interactions. SMD shows that both drugs attain different F_max_ at different time points. Reaction coordinates achieved from SMD were further used for umbrella sampling. Umbrella sampling allowed us to calculate the potential mean force (PMF) required to separate the ligand from FemX. The snapshots which were obtained from SMD trajectories were exploited for umbrella sampling. The PMF result showed that Lumacaftor required more energy than Olaparib to dislocate it from the ligand-binding site (Figure 4). Lumacaftor exhibited a higher free energy barrier at approximately 87.41 kJ/mol compared to Olaparib, which possessed a PMF of 32.31 kJ/mol. These results from umbrella sampling are completely resonating with the SMD study. These drugs were used for further experimental investigation.

**Figure 3.**
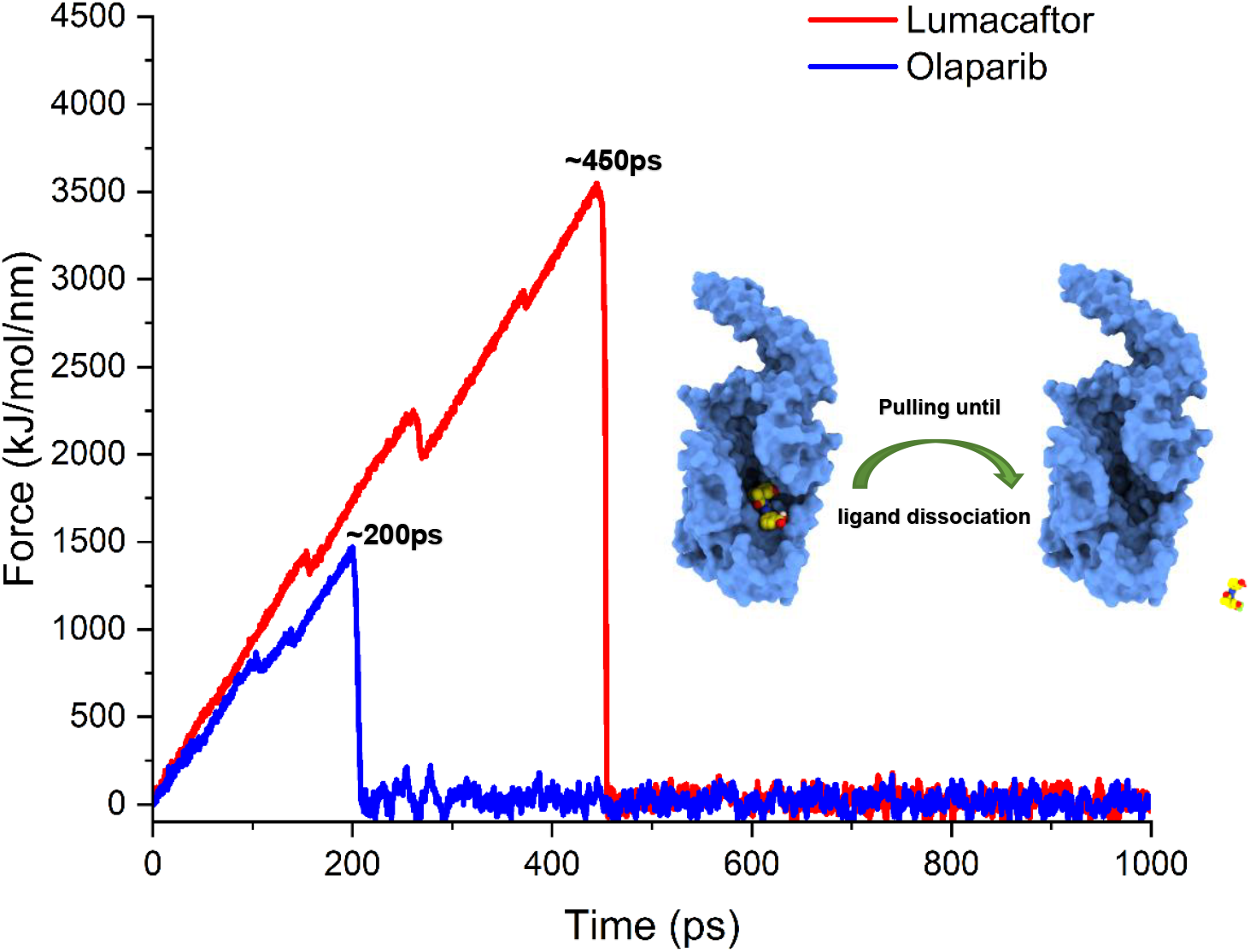
Steered molecular dynamics simulation of Lumacaftor and Olaparib showing the rupture force profile. The x-axis denotes the simulation time, whereas the y-axis denotes the pulling-out force required to dislocate the ligands from their respective binding sites.

**Figure 4.**
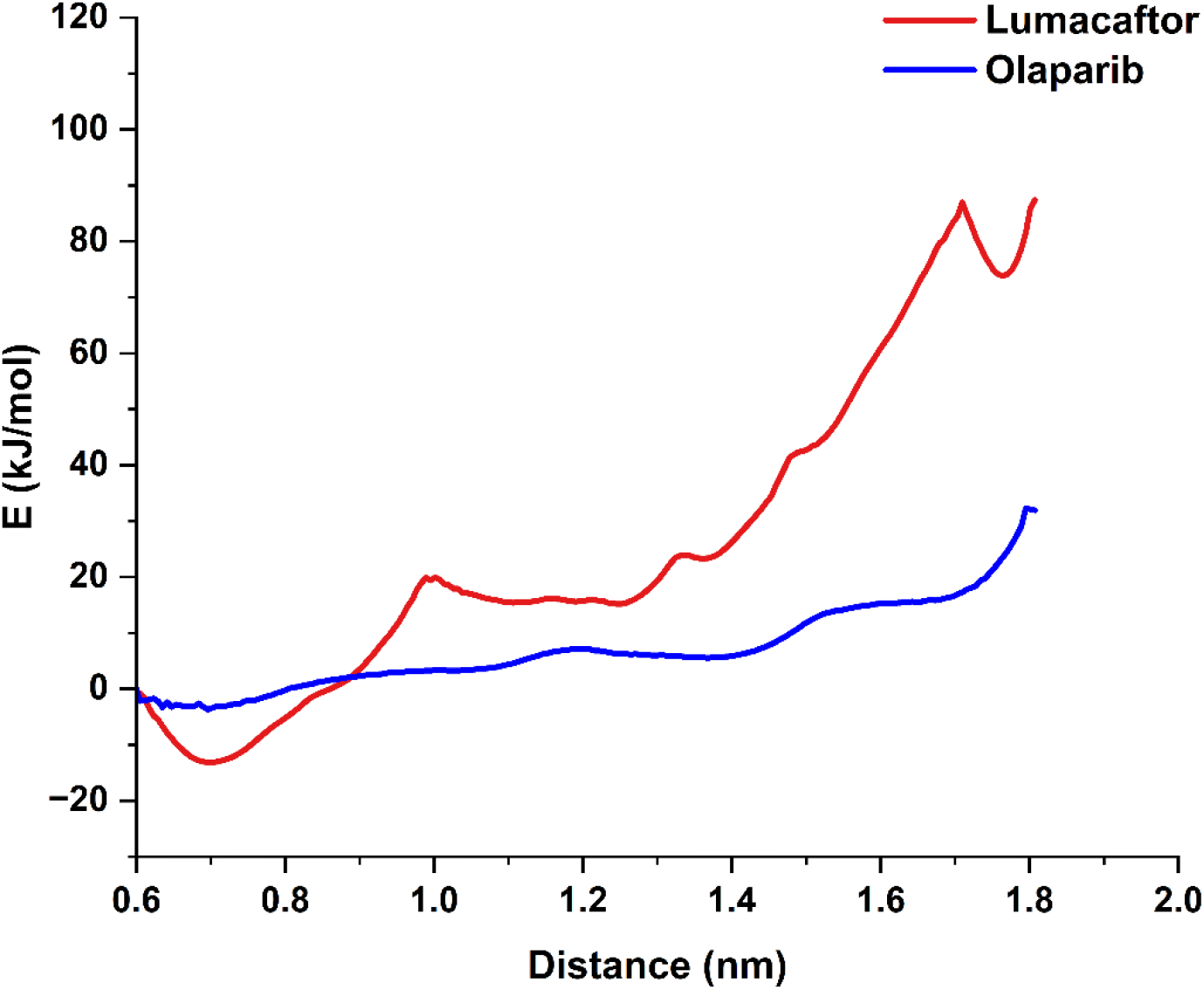
Potential mean force curves of Lumacaftor and Olaparib complexes obtained from umbrella sampling.

### 3.2 Minimum inhibitory concentration and IC_50_

Minimum inhibitory concentration was determined to assess the antibacterial activity of Lumacaftor and Olaparib against gram-positive *S.aureus*. Lumacaftor displayed antibacterial activity at an inhibitory concentration of 128 μg/mL for both the *S. aureus* strains (Figure 5). In contrast, Olaparib did not exhibit any notable antibacterial activity against *S. aureus* at any concentration tested (Supplementary Figure 1). The MIC value for Lumacaftor was determined with conc. 128 μg/mL and IC_50_ value were determined with conc. ∼65 μg/mL. These data suggested Lumacaftor has moderate inhibitory activity against *S. aureus* NCTC 8325 and MTCC 3160. These results suggest that Lumacaftor could be a promising lead compound for developing new treatments against planktonic *S. aureus*. Overall, these findings demonstrate the potential of Lumacaftor as a candidate for further exploration in the field of antibacterial drug discovery.

**Figure 5.**
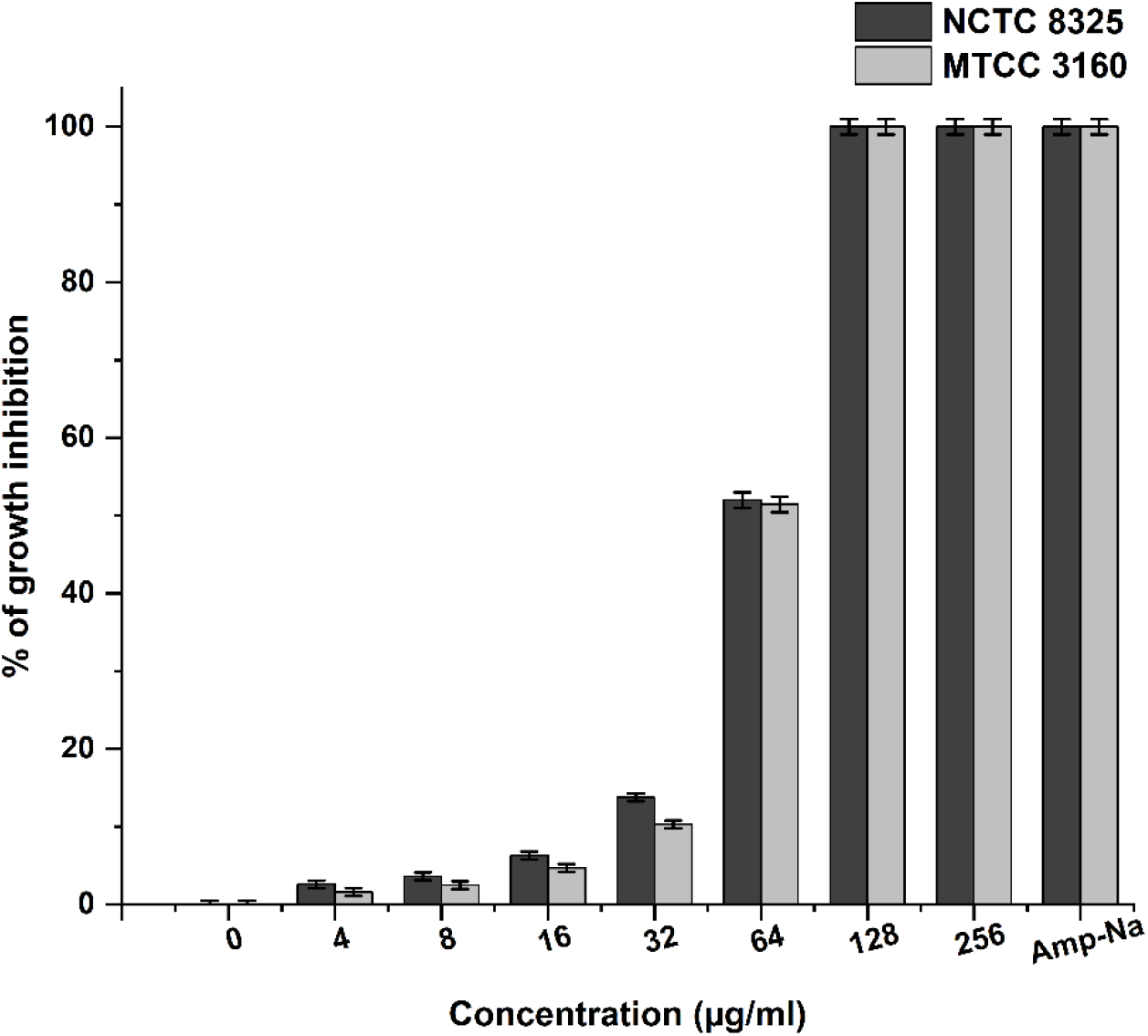
Bactericidal activity of Lumacaftor by microdilution method in terms of minimum inhibitory concentration. The MIC value for Lumacaftor was determined with a concentration of 128 μg/mL and IC_50_ value with a ∼65 μg/mL concentration for *S. aureus* NCTC 8325 and MTCC 3160 strains.

### 3.3 Biofilm inhibition

The crystal violet assay was performed to evaluate the effect of Lumacaftor and Olaparib on biofilm biomass. The results showed that Lumacaftor had a moderate impact on biofilm biomass at sub-inhibitory concentrations and a high impact on biofilm growth at the minimum inhibitory concentration after 16 hours of treatment. At sub-inhibitory concentrations, Lumacaftor was able to reduce biofilm biomass by 10-60%, while the positive control trans-chalcone was only able to achieve a ∼40% reduction (Figure 5). These data indicated the potential of Lumacaftor to inhibit staphylococcal biofilm growth. On the other hand, Olaparib did not show any biofilm inhibition activity (Supplementary Figure 2). The crystal violet assay results provided valuable insights into the potential of Lumacaftor in inhibiting biofilm growth, which can have important implications for the development of new therapies for *S.aureus*-mediated infections.

## 3. ADMET

*In-silico* ADME study analyzed the physicochemical properties, lipophilicity, water solubility pharmacokinetics, drug-likeness and medicinal chemistry properties of Lumacaftor and Olaparib. SwissADME showed Lumacaftor and Olaparib were not violating any of the Lipinski rules of five. They were found to follow the Ghose, Veber, Egan and Muegge rules with a good bioavailability score of 0.56 and 0.55 for Lumacaftor and Olaparib, respectively. The bioavailability radar (Spider plot) of Lumacaftor showed that all the 6 parameters, such as lipophilicity (XLOGP3 = 4.40), size (MV = 434.46 g/mol), polarity (TPSA = 97.75 A^2^), insolubility (LOG S{ESOL}= -5.45) and flexibility (number of rotatable bonds = 6) lie within the acceptable range except insaturation (Fraction Csp3 = 0.21; where range lies between 0.25 - 1.00) which was found to be just outside the acceptable range (Table 1) (Figure 7). Besides, the spider plot of Olaparib exhibited that all these parameters, such as lipophilicity (XLOGP3 = 1.90), size (MV = 452.41 g/mol), polarity (TPSA = 86.37 A^2^), insolubility (LOG S{ESOL}= -3.70), flexibility (number of rotatable bonds = 6) and insaturation (Fraction Csp3 = 0.33) lies within the acceptable range. ADME analysis depicted that both Lumacaftor and Olaparib have high gastrointestinal absorption (78.71% and 91.92%, respectively) and skin permeability. The BOILED-Egg model of Lumacaftor and Olaparib lies in the white region, suggesting that the two compounds are highly likely to be absorbed by the gastrointestinal tract (Figure 7). Importantly, neither drug was not crossing the blood-brain barrier (BBB), which is considered an important property to consider these drugs for usage. The pkCSM tool provided several other pieces of information about the toxicity of the drugs (Table 2). Lumacaftor and Olaparib exhibited a maximum tolerated dose of 0.824 log mg/kg/day and 0.204 (log mg/kg/day) for humans. Besides, Lumacaftor showed oral acute toxicity (LD_50_) of 2.797 (mol/kg) and oral chronic toxicity (LOAEL) of 1.424 (log mg/kg_bw/day). Olaparib showed oral acute toxicity (LD_50_) of 2.623 (mol/kg) and oral chronic toxicity (LOAEL) of 1.799 (log mg/kg_bw/day). Both drugs have no skin sensitization effect. Lumacaftor was not an inhibitor for CYP1A2, CYP2C19, CYP2D6 or CYP3A4 except CYP2C9, and Olaparib was found to be a CYP2C19 and CYP2C9 inhibitor. There were fewer chances of unwanted adverse effects due to the lower clearance and accumulation of these drugs.

**Table 1.** Pharmacokinetic evaluation of Lumacaftor using SwissADME.

| Parameters | Properties |  |
| --- | --- | --- |
|  | Lumacaftor | Olaparib |
| Molecular weight | 452.41 g/mol | 434.46 |
| Lipophilicity | 4.40 | 1.90 |
| Polarity | 97.75 Å<sup>2</sup> | 86.37 Å<sup>2</sup> |
| Insolubility | -5.45 | -3.70 |
| Flexibility in terms of the number of rotatable bonds | 6 | 6 |
| Insaturation | 0.21 | 0.33 |
| GI absorption | High | High |
| BBB permeant | No | No |
| Log Kp (Skin permeation) | -5.91 cm/s | -7.60 cm/s |

**Table 2.** Toxicology evaluation of Lumacaftor using pkCSM.

| Parameters | Properties |  |
| --- | --- | --- |
|  | Lumacaftor | Olaparib |
| Intestinal absorption (human) | 78.71% | 91.92% |
| CYP1A2 inhibitor | No | No |
| CYP2C19 inhibitor | No | Yes |
| CYP2C9 inhibitor | Yes | Yes |
| CYP2D6 inhibitor | No | No |
| CYP3A4 inhibitor | No | Yes |
| Renal OCT2 substrate | No | No |
| Maximum tolerated dose (human) | 0.824 (log mg/kg/day) | 0.204 (log mg/kg/day) |
| hERG I inhibitor | No | No |
| hERG II inhibitor | No | Yes |
| Oral Rat Acute Toxicity (LD<sub>50</sub>) | 2.797 (mol/kg) | 2.623 (mol/kg) |
| Oral Rat Chronic Toxicity (LOAEL) | 1.424 (log mg/kg_bw/day) | 1.799 (log mg/kg_bw/day) |
| Skin sensitization | No | No |

## 4. Discussion

Drug development is a time taking, costly and challenging process with a high degree of uncertainty that a drug will actually become effective or not. Here drug repurposing comes into play, which involves exploring new therapeutic usage of existing approved, discontinued, shelved and investigational drugs. Drug repurposing is a novel way of finding new uses outside the scope and offers reduced cost, Faster development timeline and regulatory approval as these drugs already have positive preclinical and safety data. This study focuses on eliminating *S. aureus* with existing market-available drugs.

Our previous work engaging virtual screening, docking, conventional molecular dynamics (CMD) and hybrid quantum mechanics/molecular mechanics (QM/MM) studies reported that Lumacaftor and Olaparib could be used as repurposing drugs against staphylococcal FemX ^18^. The docking study showed that Lumacaftor has a higher affinity in terms of K_d_ value than Olaparib. Besides, the molecular dynamics study reported that the FemX-Lumacaftor complex is more stable than the FemX-Olaparib complex. Further, molecular mechanics generalized-born surface area (MM/GBSA) calculations also indicated that Lumacaftor has higher binding energies towards FemX in comparison with Olaparib. Steered molecular dynamics is a modern method to study the ligand-receptor unbinding mechanisms ^28,29^. This current study uses SMD and umbrella sampling to estimate the pulling force and potential mean force required to dislocate the drugs from their ligand-binding sites in FemX. Lumacaftor possesses a higher pulling force and PMF than Olaparib when ligands are pulled with a time-dependent external force (Figure 3, Figure 4). The reason behind this significant variation in F_max_ peaks for the selected hits is due to the different interaction profiles of the drugs with FemX. Previous *in-silico* and *in-vitro* studies also reported that drugs possess the same dissociation trend during SMD and umbrella sampling ^28–33^. Lumacaftor and Olaparib possess two and three H-bonds with FemX, respectively. Interaction between Lumacaftor and FemX is mainly stabilized by strong π−π interactions between aromatic residues of receptor and aromatic rings of ligands ^18^. SMD and umbrella sampling data support the findings from conventional molecular dynamics and hybrid QM/MM studies, which in turn offer the acceptability of the current study.

The peptidoglycan layer maintains the cell-membrane integrity and is essential for bacterial survival. The peptidoglycan biosynthesis pathway is the most widely targeted area for designing new and potent antibiotics ^34,35^. Comprehensive information on the mechanism of methicillin resistance paved the way for discovering auxiliary factors such as Fem, which regulates the methicillin resistance in *S.aureus*. FemXAB activity is highly substrate-specific, where FemX mutations/deletion is lethal to *S. aureus* ^16^. Ampicillin-sodium, which is used in this study as a positive control, is a member of the extended-spectrum β-lactam family. Nowadays, resistance to β-lactams in *S.aureus*-mediated infections is a serious healthcare concern ^36^. The prevalence of antibiotic resistance among pathogens is a growing problem that draws attention to the development of new antibiotics ^37^. However, β-lactam antibiotics cause medication error that has been recognized as a common and serious threat to patient safety ^38^. β-lactam antibiotics are found to be involved with severe adverse effects ^39,40^. Thereby, new drugs are being explored against *S. aureus* using a drug-repurposing approach ^41–43^. Our study showed that Lumacaftor has decent bactericidal activity (MIC: 128 μg/mL), but Olaparib had no potential against *S. aureus* MTCC 3160 and NCTC 8325 strains (Figure 5). Surprisingly, we have found that Lumacaftor can inhibit biofilm formation 10-60% with the sub-inhibitory concentrations, which indicates its potential as an anti-biofilm agent. Previous studies also showed that drugs could significantly inhibit biofilm formation at sub-inhibitory concentrations ^44,45^. CMD and SMD studies also indicated that Lumacaftor might have more potential against *S. aureus*. Lumacaftor was significantly able to inhibit staphylococcal biofilm formation (Figure 6). Previous studies reported that FDA-approved drugs with higher F_max_ values possess higher biological activity ^46,47^. These previous reports evaluate the robustness of our currently employed approach. Based on the literature, no such standard drug is available against staphylococcal FemX, so we could not perform any comparison. Combination antibiotic treatment against bacterial infections is an attractive alternative as it could address the shortcomings of most antibiotics. Recently, antibiotic combinations have been used against *S. aureus* and other bacteria to increase the efficiency and reduce the resistance to any individual drugs ^48–51^. Thereby, we suggest that the synergistic use of Lumacaftor with other antibiotics can increase drug activity and bacterial elimination.

**Figure 6.**
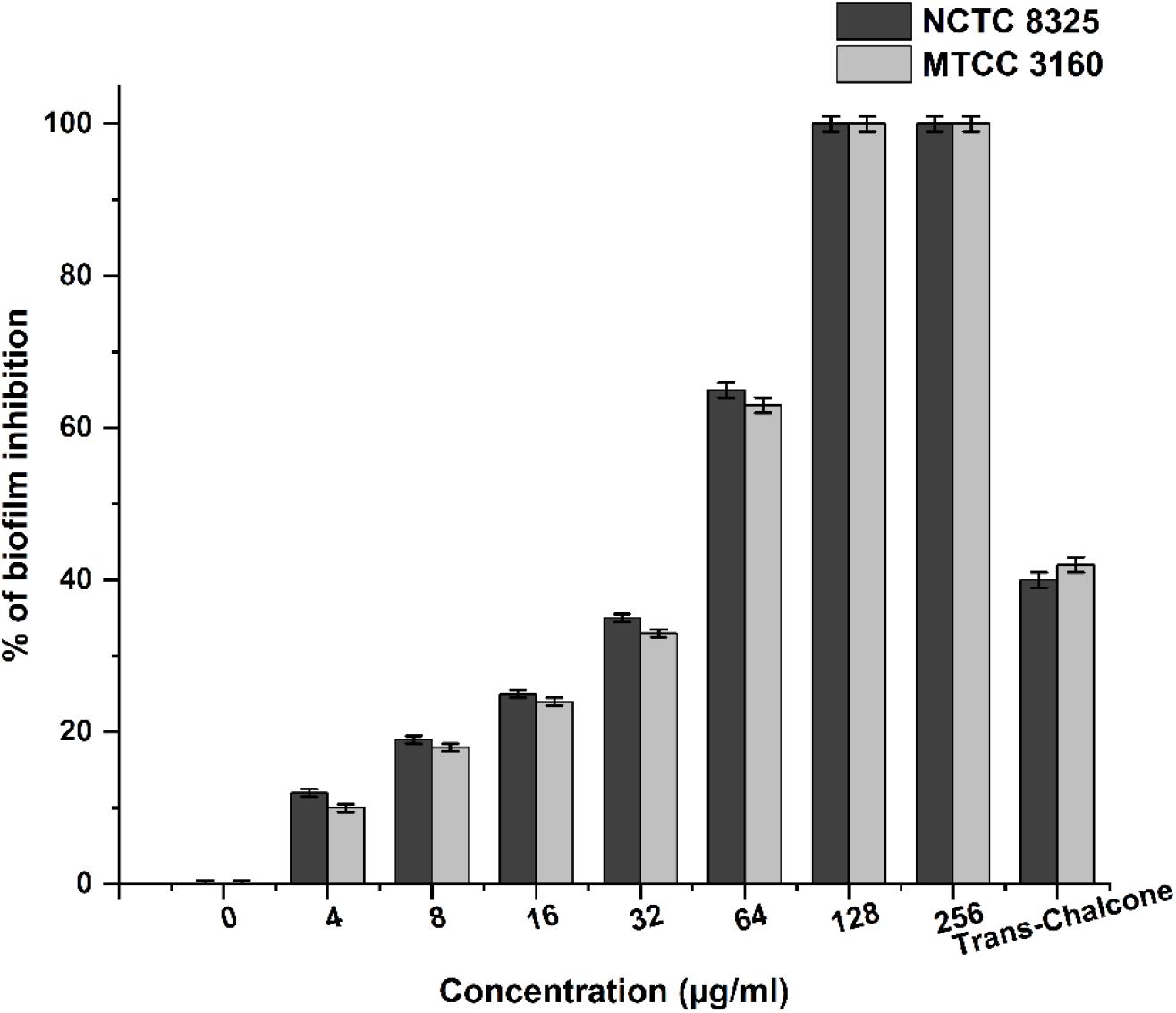
Susceptibility of *Staphylococcus aureus* biofilms to inhibition by Lumacaftor. The percentage of biofilm inhibition after treatment with Lumacaftor indicates the efficacy of the drug against *S. aureus* NCTC 8325 and MTCC 3160 strains.

Many medications have failed during the drug development process due to poor pharmacokinetics and toxicity issues ^52^. Problems that arise during the drug development process could be resolved early on. *In-silico* ADMET methods are the first step in this pipeline process to assess the issues of new chemical compounds ^53^. ADMET factors reveal how chemical substances behave within a living being. These techniques save time by avoiding lead candidates that would be harmful or would be converted by the body into an inactive form. These ADMET parameters disclose the behaviour of the chemical compounds in a living organism. In this study, Lumacaftor and Olaparib were evaluated for their pharmacokinetic potential. SwissADME and pkCSM use a combination of in-silico models and experimental data to make their predictions using the input of accurate and complete molecular information such as 2D/3D structure, charges, and tautomeric forms ^26,27^. The BOILED-egg model predicts the gastrointestinal absorption and blood-brain barrier permeation of the given drug ^54^. Both drugs show high GI absorption values with no BBB permeation potential, indicating their acceptability as promising therapeutic agents (Figure 7). The Lipinski filter is the pioneer rule of five, and none of the drugs is found to violate any of the rules. Besides, the cytochrome P450 (CYP) superfamily of isoenzymes is crucial in drug elimination through metabolic pathways. Inhibition of five major CYP isoenzymes such as CYP1A2, CYP2C19, CYP2CP, CYP2D6 and CYP3A4, is certainly one major cause of pharmacokinetics-related drug-drug interactions that may lead to toxic effects due to accumulation and lower clearance of the drug ^55–57^. Lumacaftor was found to be a CYP2C9 inhibitor, whereas, Olaparib was found to be a CYP2C19 and CYP2C9 inhibitor (Table 2). These data indicated that Lumacaftor could be less toxic than Olaparib. Moreover, these are FDA-approved drugs, which means they have already been validated for therapeutic use, outweighing the intended use risks. And importantly, none of the drugs can elicit any allergic response in susceptible individuals due to a lack of skin sensitization effect. The collective data suggest that Lumacaftor may be considered as a lead molecule or a prototype for developing new derivatives.

**Figure 7.**
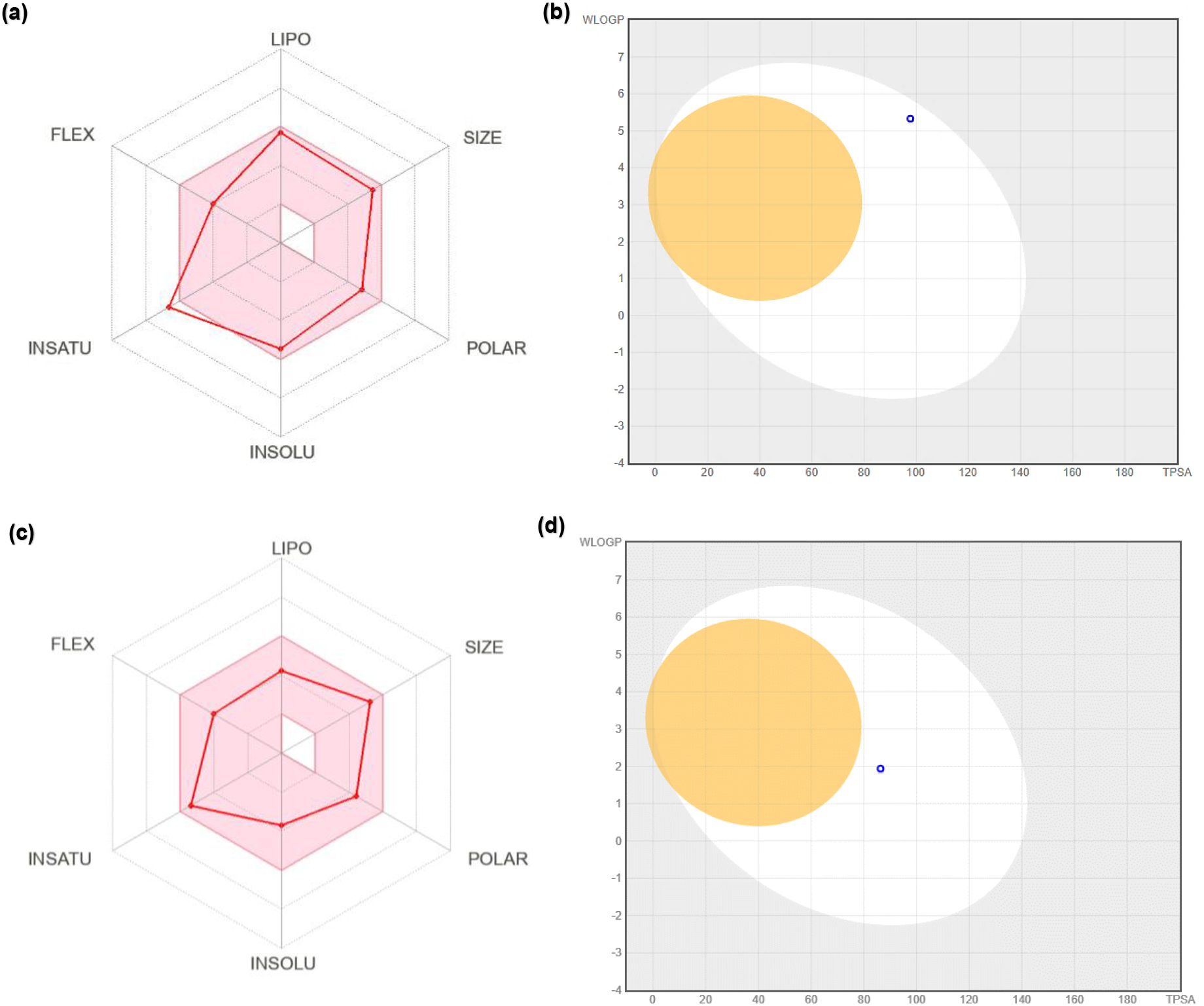
Bioavailability radar and BOILED-egg models of Lumacaftor and Olaparib based on physicochemical descriptors. The pink zone in the bioavailability radar is the ideal physicochemical space for oral bioavailability in the case of Lumacaftor (a) and Olaparib (c). BOILED-egg models predicted the GI absorption and blood-brain barrier permeation of Lumacaftor (b) and Olaparib (d).

## Conclusion

This study evaluates the activity of two drugs, Lumacaftor and Olaparib, against *Staphylococcus aureus*. The study targets *S. aureus* FemX - the protein involved in building the bacterial cell wall. *In-silico* analysis shows that Lumacaftor requires significantly more energy (potential mean force) than Olaparib to dislodge from the binding site of FemX. Lumacaftor has moderate bactericidal activity with a MIC value of 128 μg/mL and inhibits the planktonic growth of the S. aureus MTCC3160 and NCTC 8325 strains. On the other hand, Olaparib does not show any bactericidal activity. Moreover, Lumacaftor inhibits biofilm formation when treated with sub-inhibitory concentration, but Olaparib does not inhibit it. It may be concluded that Lumacaftor can be a potential lead molecule against *S. aureus* for further development as an antibiotic.

## Data and Software Availability

GROMACS 2020.0 is an open-source package freely available from the source (http://ftp.gromacs.org/pub/gromacs/gromacs-2020.tar.gz). Protein structure is available in Protein Data Bank (PDB ID: 6SNR). Ligand information is publicly available in the PubChem database (https://pubchem.ncbi.nlm.nih.gov/).

## Author contribution

SR conceptualized the study, performed steered molecular dynamics simulation and umbrella sampling experiments and wrote the draft manuscript. SN performed MIC and biofilm inhibition studies. UM and AKD supervised the work and improvised the final manuscript.

## Acknowledgements

This work has been carried out with financial assistance from the Science and Engineering Research Board, Government of India (GOI), File no. CRG/2020/002622. SR acknowledges the Indian Institute of Technology Kharagpur for individual fellowship. The authors would like to acknowledge Ranabir Majumder and Rituparna Saha, IIT Kharagpur, for critical suggestions. This work used the computational resources of the PARAM shakti supercomputing facility of IIT Kharagpur (www.hpc.iitkgp.ac.in), established under the National Supercomputing Mission of the Government of India (GoI) and supported by CDAC, Pune.

