## Supplementary Figure 1, Supplementary Figure 2 for "Targeting staphylococcal cell-wall biosynthesis protein FemX through steered molecular dynamics and drug-repurposing approach"

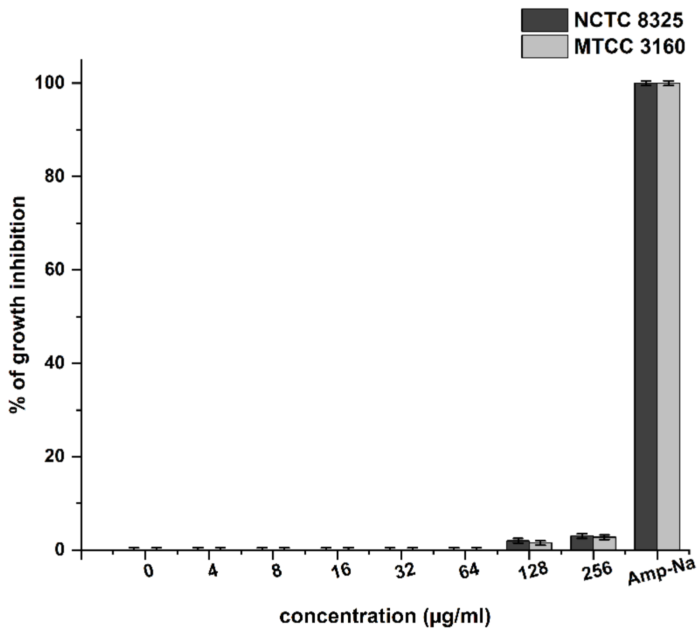


**Supplementary Figure 1. Bactericidal activity of Olaparib by microdilution method in terms of minimum inhibitory concentration. Olaparib did not affect the growth of *S. aureus* NCTC 8325 and MTCC 3160 strains.**

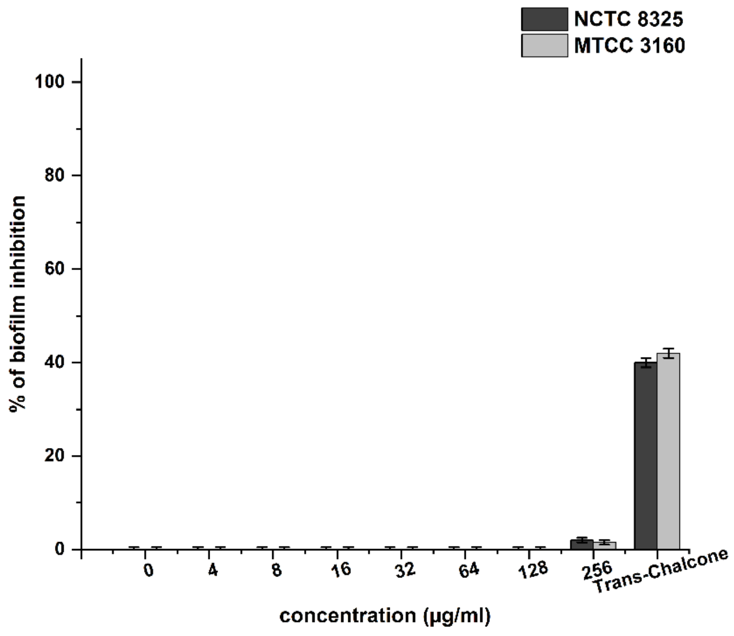


**Supplementary Figure 2. The percentage of biofilm inhibition after treatment with Olaparib indicates the efficacy of the drug against *S. aureus* NCTC 8325 and MTCC 3160 strains. Olaparib could not inhibit the biofilm formation of *S. aureus* strains.**

**Supplementary Table 1. Biofilm eradication by Lumacaftor in *S. aureus* strains.**

| **Lumacaftor (µg/mL)** | **Biofilm eradication % *in S. aureus* MTCC 3160** | **Biofilm eradication % in *S. aureus* NCTC 8325** |
|---|---|---|
| 0 | 0 | 0 |
| 4 | 0 | 0 |
| 8 | 0 | 0 |
| 16 | 0 | 0 |
| 32 | 0 | 0 |
| 64 | 0 | 0 |
| 128 | 0.6 | 0.8 |
| 256 | 1.60 | 1.92 |
| Amp-Na (6.25 µg/ml) | 1.50 | 1.45 |
| Amp-Na (160 µg/ml) | 50.94 | 54.50 |

**Supplementary Table 2. Biofilm eradication by Olaparib in *S. aureus* strains.**

| **Olaparib (µg/mL)** | **Biofilm eradication % *in S. aureus* MTCC 3160** | **Biofilm eradication % in *S. aureus* NCTC 8325** |
|---|---|---|
| 0 | 0 | 0 |
| 4 | 0 | 0 |
| 8 | 0 | 0 |
| 16 | 0 | 0 |
| 32 | 0 | 0 |
| 64 | 0 | 0 |
| 128 | 0 | 0 |
| 256 | 0 | 0 |
| Amp-Na (6.25 µg/ml) | 1.50 | 1.45 |
| Amp-Na (160 µg/ml) | 50.94 | 54.50 |
